# The evolution of ageing in cooperative breeders

**DOI:** 10.1101/2022.03.04.482977

**Authors:** Jan J. Kreider, Boris H. Kramer, Jan Komdeur, Ido Pen

## Abstract

Cooperatively breeding animals live longer than their solitary counterparts. The traditional explanation for this is that cooperative breeding evolves more readily in long-lived species. Here, we reverse this argument and show that long lifespans are an evolutionary consequence of cooperative breeding. Natural selection favours a delayed onset of senescence in cooperative breeders, relative to solitary breeders, because cooperative breeders have a delayed age of first reproduction due to reproductive queueing. Especially long lifespans evolve in cooperative breeders with age-dependent reproductive queueing. Finally, we show that lower genetic relatedness among group members leads to the evolution of longer lifespans. This is because selection against higher mortality is weaker when mortality reduces competition between relatives. Our results link the evolutionary theory of ageing with kin selection theory, demonstrating that the evolution of ageing in cooperative breeders is driven by the timing of reproduction and kin structure within breeding territories.

## Introduction

The evolution of sociality is associated with changes in life history, especially lifespan^1–3^. As demonstrated in birds^4,5^ and mole rats^6,7^ (though not in mammals in general^8,9^), cooperatively breeding species often have longer lifespans than solitary species, and the reproductive castes of eusocial insects outlive solitary insects by several orders of magnitude^10^. The prevalence of long lifespans in cooperative breeders has been interpreted as evidence in favour of the so-called “life history hypothesis” of cooperative breeding^11–14^. This hypothesis posits that particular life history traits, such as low adult mortality, facilitate the evolution of cooperative breeding because lower mortality reduces the rate of territory turnover. If access to breeding territories is restricted, it can be beneficial for individuals to remain near their natal territory and help raise offspring of relatives (“indirect fitness benefits”)^15–19^ and/or to queue for a breeding territory with a chance to inherit a breeding position (“direct fitness benefits”), even if this requires alloparental care towards non-relatives^20–23^. Thus, the logic of the “life history hypothesis” implies that cooperative breeders are long-lived because the longevity of their solitary ancestors played a causal role in the evolution of cooperative breeding.

However, the logic may also be reversed – rather than being a cause of cooperative breeding, long lifespans could be a consequence of it. This seems to follow from Hamilton’s classical theory on the evolution of senescence^24^, which demonstrated that the strength of natural selection against higher mortality is maximal and constant until the age of first reproduction and declines with age afterwards. Consequently, a delayed age of first reproduction implies a delayed onset of senescence and the evolution of longer lifespans. In cooperative breeders, sexually mature helpers typically have to wait for an extended period in a reproductive queue and therefore have a delayed age of first reproduction^5^. As a result, cooperatively breeding species should evolve longer lifespans than otherwise similar solitary species. However, Hamilton’s model did not explicitly consider effects of sociality^25^, hence the need for a formal model of the evolution of ageing in cooperative breeders.

Here, we present an evolutionary individual-based simulation model to derive predictions for the evolution of ageing in cooperatively breeding organisms. The model represents a population of individuals whose lifespans evolve due to the accumulation of mutations with age-specific effects on survival, as in Medawar’s mutation accumulation theory of ageing^26^. We simulated the evolution of ageing both in solitarily and cooperatively breeding organisms, representing a broad range of biological systems (Fig. 1). As the productivity benefits of helpers, i.e. the increase in the reproductive output of the dominant breeders caused by the presence of helpers, can vary between cooperatively breeding species, we investigated how such productivity benefits affect the evolution of ageing. In cooperative breeders, reproductive queues may consist of highly related individuals but also of non-related individuals. Therefore, we evaluated the effect of kin structure within breeding territories on the evolution of ageing in cooperative breeders.

**Fig 1.**
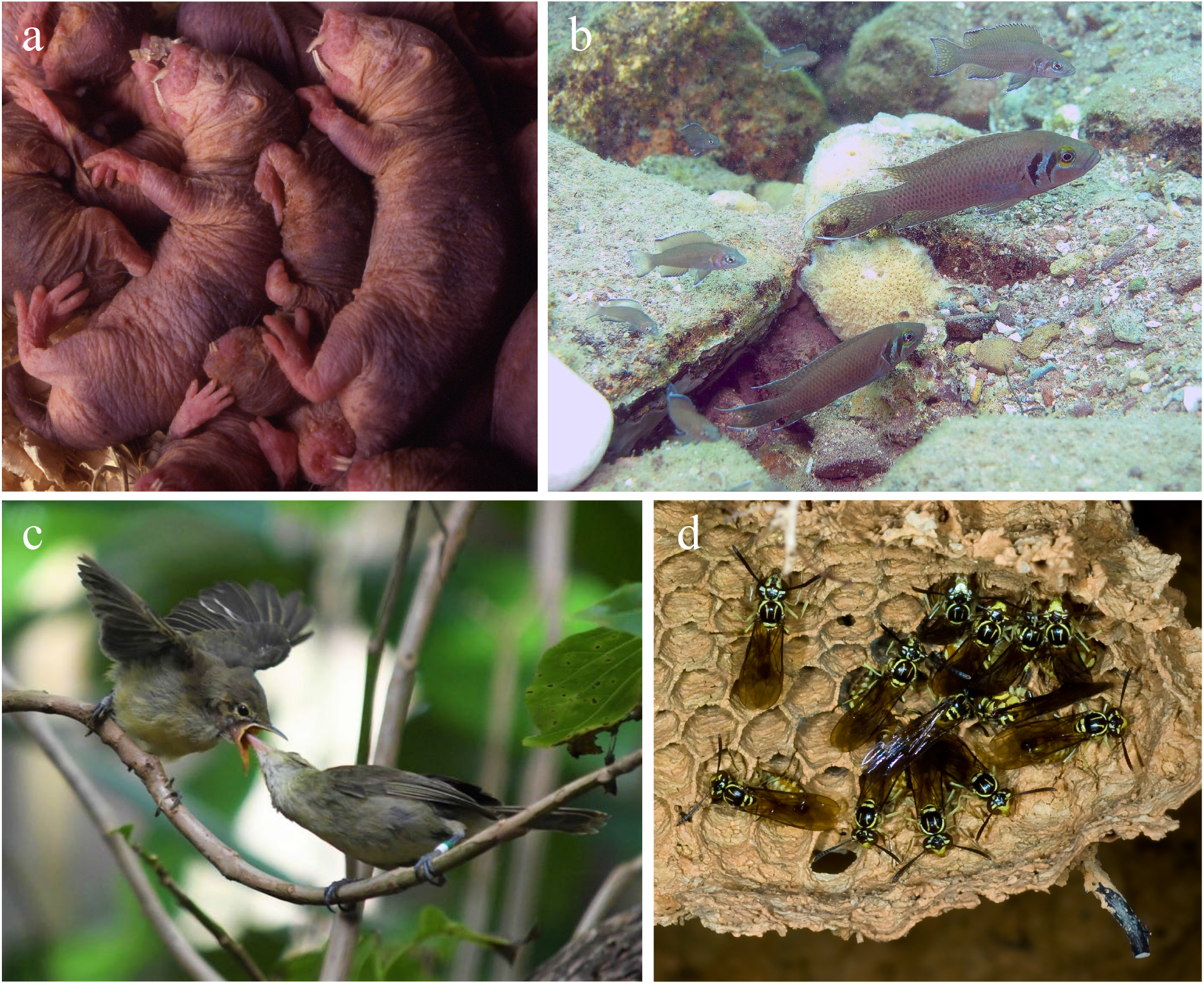
Cooperatively breeding organisms from disparate phylogenetic lineages. **(a)** The naked mole rat (*Heterocephalus glaber*) is a long-lived rodent that lives in closely-related family groups and exhibits age-dependent reproductive queueing^27–30^ (copyright: Chris Faulkes). **(b)** The Lake Tanganyika princess (*Neolamprologus pulcher*) is a cooperatively breeding cichlid with low relatedness within groups and size-dependent reproductive queueing^31–33^ (copyright: Dario Josi). **(c)** The Seychelles warbler (*Acrocephalus sechellensis*) has low within-group relatedness and age-independent reproductive queueing^34–36^ (copyright: Charli Davies). **(d)** The tropical hover wasp (*Liostenogaster flavolineata*) has high relatedness within groups and age-dependent reproductive queueing^37–40^ (copyright: David Baracchi).

## Results

### Reproductive queueing causes the evolution of longer lifespans through a delay of reproduction

If non-breeding individuals can inherit a breeding territory later in their life, either through queueing for an unoccupied breeding territory globally as floaters (“solitary queueing”) or locally within a breeding territory as helpers (“cooperative breeding”), longer lifespans evolve than in the absence of queueing (“solitary”). Longer lifespans coincide with an increase in the age of first reproduction (Fig. 2). In cooperative breeders, queueing positions within breeding territories may depend on the helpers’ age relative to that of other helpers in the group (see examples in Fig. 1). Under such age-dependent queueing (“cooperative breeding weakly age-dependent” and “cooperative breeding strictly age-dependent”), longer lifespans evolve and the age of first reproduction increases relative to age-independent reproductive queueing (“cooperative breeding age-independent”). The increase of lifespans through age-dependent reproductive queueing compared to age-independent reproductive queueing is even stronger when maximum lifespans are increased from 20 to 40 (Fig. S1).

**Fig 2.**
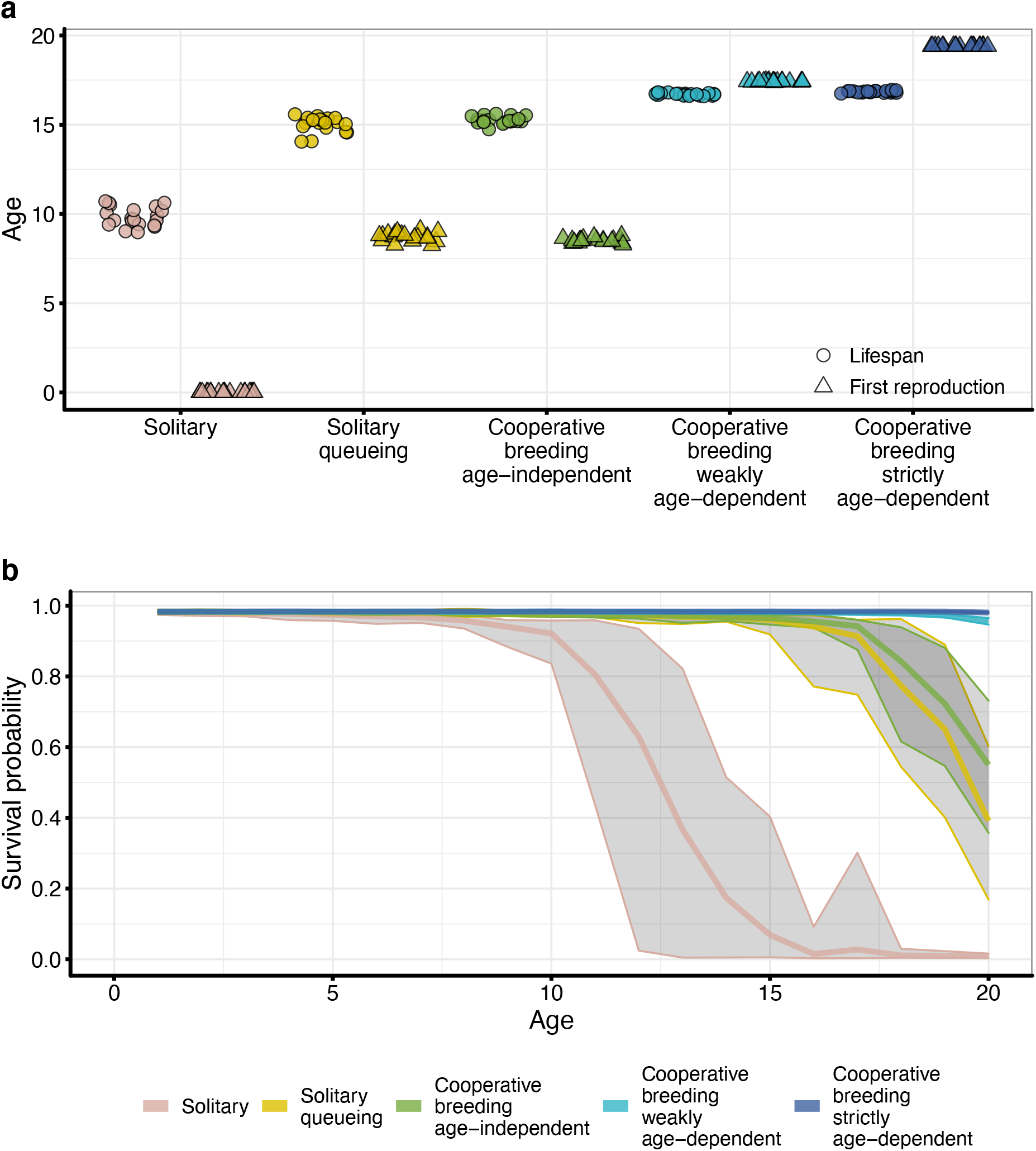
The evolution of ageing in solitary and cooperative breeders. **(a)** Evolved lifespans (circles) and age of first reproduction (triangles) for different solitary and cooperative breeding scenarios. Data points (*n* = 20) are the population mean at the end of a replicate simulation. **(b)** Age-specific survival probabilities for different solitary and cooperative breeding scenarios. Bold lines represent the mean of age-specific survival probabilities and grey shaded areas the range across replicate simulations. For survivorship curves, see Fig. S2. Parameters: *d* = 1.0 (dispersal rate), *a* = 2.5 (“maximum productivity”).

### Enhanced productivity of cooperative breeding has a weak effect on the evolution of lifespans

In cooperatively breeding species, helpers typically have a positive effect on the breeder’s reproduction or on offspring survival^41–46^. However, it has been shown that these positive helper effects diminish at large group sizes^47,48^. Therefore, we modelled reproductive output of breeders as an increasing function with diminishing returns with respect to the number of helpers present, converging towards a “maximum productivity”, the maximum number of offspring that can be produced at large group sizes (for details, see Methods). Increased “maximum productivity” leads to larger group sizes, but in all scenarios of cooperative breeding, irrespective of whether reproductive queueing is age-dependent or independent of age, “maximum productivity” has only a small effect on the evolution of lifespans (Fig. 3). This is also the case when dispersal rates change (Fig. S3).

**Fig 3.**
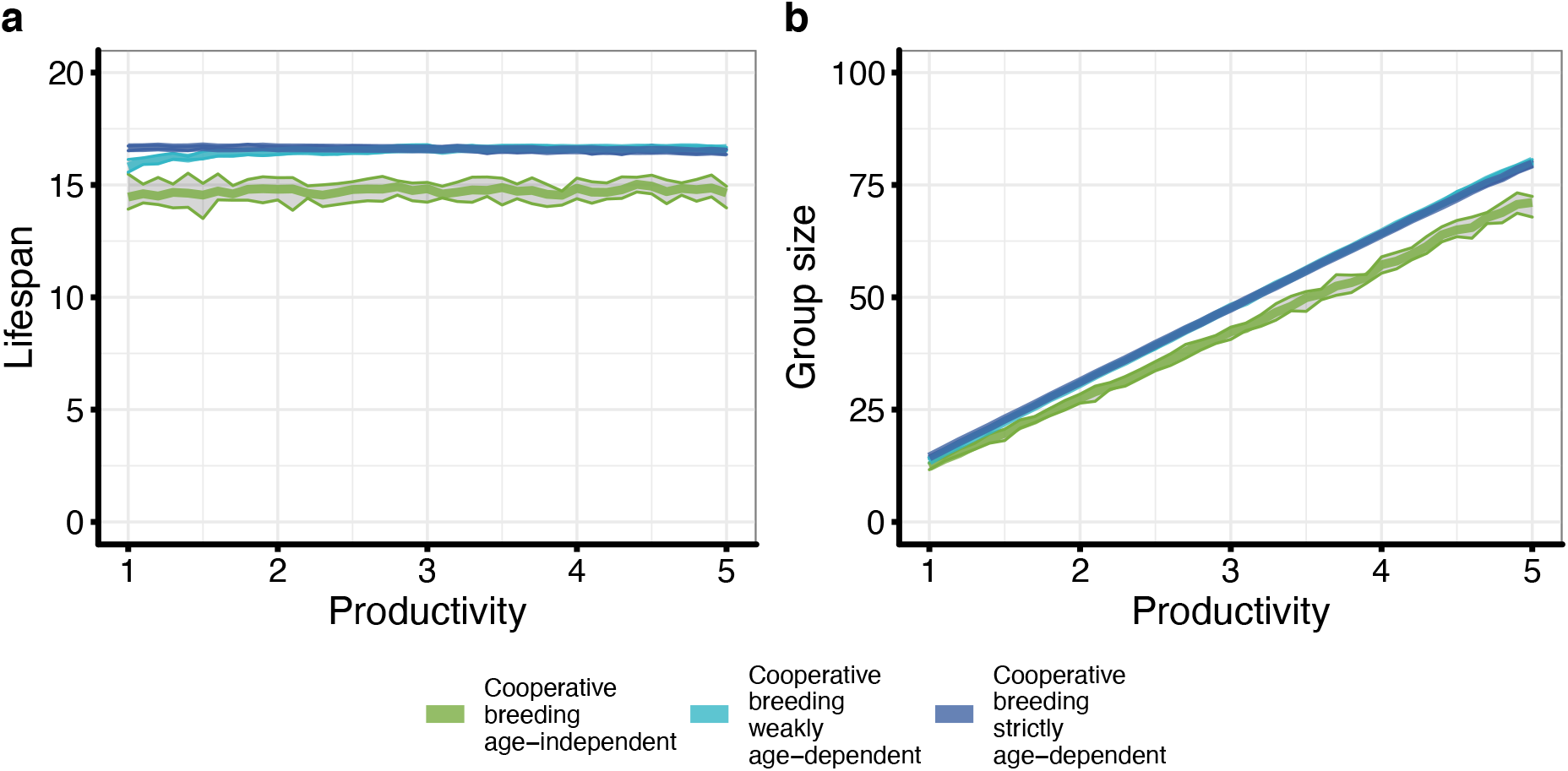
The effect of maximum productivity on the evolution of ageing in cooperative breeders. **(a)** Evolved lifespans depending on “maximum productivity” in different cooperative breeding scenarios. “Maximum productivity” is a model parameter that determines the number of offspring produced at large group sizes. **(b)** The effect of “maximum productivity” on group size. We ran each “maximum productivity” between *a* = 1.0 and *a* = 5.0 with steps of 0.1 (*n* = 10). Bold lines represent the mean evolved lifespan and grey areas the range of evolved lifespans across replicate simulations. Parameters: *d* = 0.5 (dispersal rate).

### Lower relatedness within breeding territories leads to the evolution of longer lifespans

In cooperatively breeding species, relatedness between individuals that share a breeding territory can vary (see examples in Fig. 1). We consequently allowed offspring either to be philopatric and become a helper in the natal breeding territory or to disperse and become a helper in another breeding territory. We estimated relatedness between the breeder and a random helper from the same breeding territory. Under age-independent reproductive queueing, higher dispersal rates result in lower relatedness within breeding territories and lead to the evolution of longer lifespans than under lower dispersal rates (Fig. 4). When, in contrast, reproductive queueing is age-dependent, relatedness within breeding territories is relatively low. This is because under age-dependent reproductive queueing age of first reproduction increases compared to age-independent reproductive queueing, and this increases the number of generations of females waiting in the reproductive queue, thus diluting relatedness. Lifespans are consequently not as strongly affected by dispersal rate.

**Fig 4.**
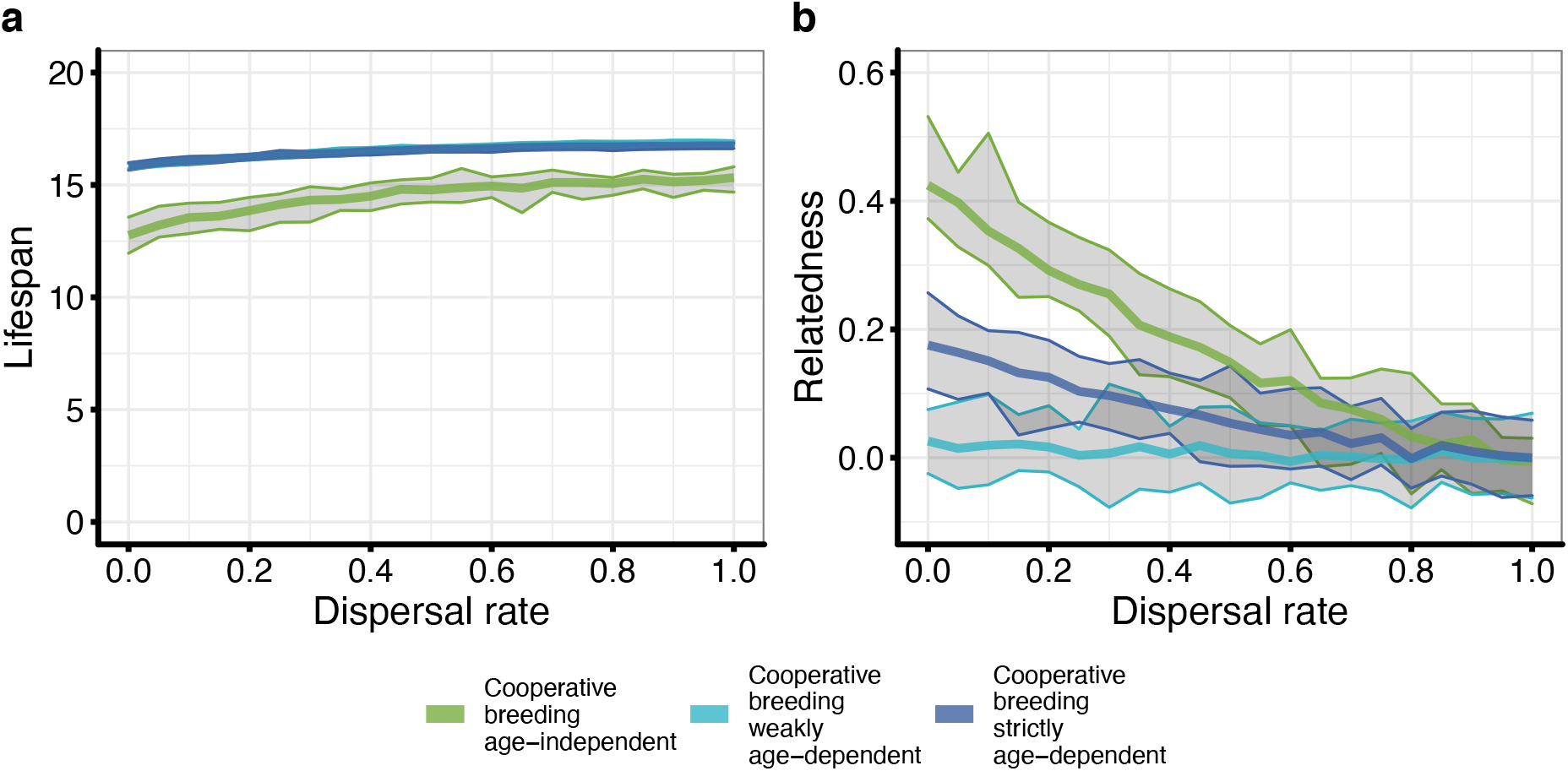
The effect of kin structure on the evolution of ageing in cooperative breeders. **(a)** The effect of dispersal rate on evolved lifespans in different cooperative breeding scenarios. **(b)** The effect of dispersal rate on relatedness between breeders and helpers in different cooperative breeding scenarios. Dispersal rates were varied between *d* = 0 and *d* = 1 in steps of 0.05 (*n* = 20). Bold lines represent the mean and grey areas the range across replicate simulations. Parameters: *a* = 2.5 (“maximum productivity”).

## Discussion

The “life history hypothesis” of cooperative breeding explains the prevalence of long lifespans in cooperative breeders by proposing that long lifespans facilitate the evolution of cooperative breeding^11–14^. Using evolutionary individual-based simulations we show that, conversely, long lifespans in cooperative breeders can also evolve as a consequence of cooperative breeding rather than being a cause for its evolution. Our results do not contradict the proposition that long lifespans facilitate the evolution of cooperative breeding – a claim which has also received formal support^12^. Indeed, it seems likely that longevity and sociality are mutually reinforcing^49–51^. However, our model also shows that evolved lifespans did not differ between solitary organisms with queueing floaters and cooperatively breeding organisms with age-independent reproductive queueing in fully outbred populations. In both cases, queueing individuals compete exclusively with unrelated individuals and their probability to achieve breeding status is independent of their age. This shows that not cooperative breeding per se, but instead the timing of first reproduction is the main determinant of the evolved onset of senescence in both solitary and cooperatively breeding organisms. In queueing systems, age at first reproduction, in turn, is influenced by lifespan as longer lifespans decrease the rate of territory turnover and thus delay the onset of reproduction. Consequently, there is positive feedback between age of first reproduction and evolved lifespans in queueing systems.

The argument that long lifespans evolve as a consequence of cooperative breeding is not unprecedented, although different mechanisms have been proposed to cause these long lifespans. For instance, recent work suggests that breeders live longer because they can afford to reduce their costly parental investment in the presence of helpers^52–54^. In our model, in contrast, longer lifespans evolve in cooperative breeders due to the delayed age of first of reproduction^24^. Both of these mechanisms are not mutually exclusive, and the simultaneous presence of both might increase lifespans even more than each on its own.

Comparative studies have yielded somewhat ambiguous findings on the association between lifespan and cooperative breeding – some found longer lifespans in cooperative breeders compared to solitary species^4–7^ and some did not^8,9^. Our results show large variation of evolved lifespans within solitarily breeding organisms, depending on whether individuals can queue to obtain a breeding territory or not, and within cooperatively breeding organisms, depending on whether reproductive queueing is age-dependent or independent of age. These factors might consequently be better predictors for interspecific lifespans differences than a dichotomous distinction between solitary and cooperative breeding.

Our model furthermore shows that positive effects of helping on reproductive output only have a minor effect at best on the evolution of lifespans in cooperatively breeding organisms, although maximum productivity clearly affected group size. In eusocial species, colony size positively correlates with the divergence of queen and worker lifespans^55^. However, in cooperative breeders there seems to be only a small effect of group size on the evolution of ageing, as our model suggests. This is probably because the productivity benefits of helping to raise related offspring are to some extent counteracted by the simultaneous increased competition for breeder positions^56^.

In contrast to the lack of productivity effects, the presence of relatives within breeding territories did affect the evolution of lifespans. Higher within-territory relatedness – due to greater offspring philopatry – decreased evolved lifespans because selection against higher mortality is weaker if mortality reduces competition between relatives. It has been suggested that indirect fitness benefits that are gained post-reproductively facilitate the evolution of extended post-menopausal lifespans, as found in humans and some species of whales, and thus higher relatedness between group members should lead to the evolution of longer lifespans^57–60^. However, in cooperative breeders, indirect fitness benefits can also be gained pre-reproductively, because an individual’s death reduces the waiting time for its relatives behind it in the queue, and this leads to a pre-reproductive decline of the age-specific force of selection. In line with this argumentation, relatively shorter lifespans evolved in our simulations when relatedness within breeding territories was high. Consistent with this result, a comparative study found that species-specific survival in cooperatively breeding birds is positively correlated with species-specific promiscuity, which in turn is negatively correlated with intra-group relatedness^5^.

Lastly, age of first reproduction increases in some human societies^61^, despite a concomitant decrease in the age at sexual maturity^62^. At the same time, human lifespans have increased dramatically over past decades^63^. This trend is probably not caused by evolutionary adaption but by reduced mortality due to, for instance, improved medical care and nutrition^64^. However, our model also predicts that the delay of reproduction, as observed in some human societies, should impose stronger selection against pre-reproductive mortality, and thus lead to an increase of human lifespans.

Overall, our model makes an important link between the evolutionary theory of ageing and kin selection theory, demonstrating that timing of reproduction and kin structure are the most important drivers for the evolution of ageing in cooperative breeders.

## Methods

### Solitary and cooperative breeding scenarios

We developed an evolutionary individual-based simulation model. The model represents a population with a fixed number of breeding territories *N* (for all model parameters and their default values, see Table 1). Each breeding territory is initialized with one breeding female. We modelled five different breeding systems: (1) *Solitary breeding:* Females disperse at independence and compete for empty breeding territories. Females that fail to obtain a breeding territory die during that breeding season. (2) *Solitary breeding with queueing:* Females disperse at independence to become floaters that “queue” globally for breeding territories. (3) *Cooperative breeding with age-independent queueing:* Females become helpers who queue locally in a breeding territory with a chance to inherit the breeder position after the breeder’s death. A new breeder is randomly selected from all helpers in the breeding territory. (4) *Cooperative breeding with weakly age-dependent queueing:* Females queue locally in a breeding territory. Their probability to inherit the breeding position increases with their relative age in the breeding territory. (5) *Cooperative breeding with strictly age-dependent queueing:* Females again queue locally in a breeding territory. Upon a breeder’s death, the oldest helper always becomes the new breeder. In all three cooperative breeding scenarios, helpers can additionally be selected randomly to become a breeder if a breeding territory other than their own becomes empty. Furthermore, in all cooperative breeding scenarios, helpers disperse at independence with probability *d* to become a helper in another breeding territory than their natal breeding territory. We assume that only females can become helpers. Males always disperse from their natal breeding territory and mate randomly with females from the entire population.

**Table 1.**
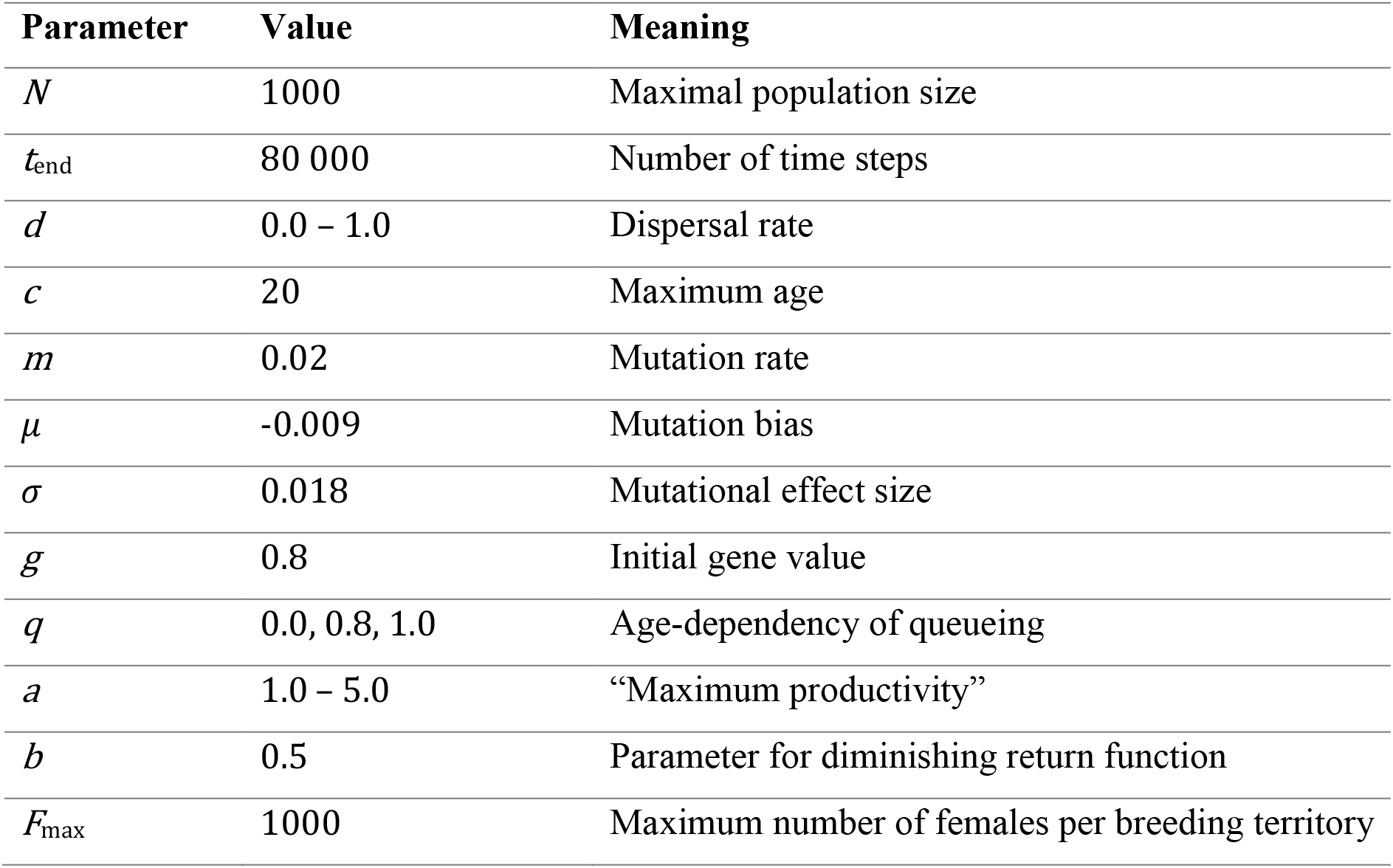
Model parameters. If parameter ranges are given, the exact parameter value is stated in the figure caption. For further parameter exploration, see the supplementary materials.

### Genetics

Individuals have diplodiploid genetics and carry genes that are associated with a number that ranges from 0.0 to 1.0. The gene values of the two homologous diploid loci are averaged to determine an age-specific survival probability. Gene values are mutated with a genome-wide mutation rate *m* at the time of emergence of a new individual. We assume that mutational effects are biased towards negative values, and thus lower survival to model accumulation of age-specific deleterious mutations. If a mutation occurs, all gene values of one randomly selected haploid gene block are separately mutated by sampling a normal distribution with a mean of *μ* < 0 (“mutation bias”) and a standard deviation of *σ* (“mutational effect size”). If a mutation causes the gene values to exceed their lower limit of 0.0 or upper limit of 1.0, they are set back to the respective limit. The haploid gene blocks can be recombined during gene transmission. We varied the default parameters for mutation rates, mutation biases and mutational effect sizes in the supplementary materials (Fig. S4). All gene values were initialized with values of *g* (see Table 1).

### Survival

Females survive or die depending on their genetically determined age-specific survival probability. Additionally, females die when they reach the maximum age *c*. We varied the default maximum age in the supplementary materials (Fig. S1, S5). For simplicity, males always die after one time step.

### Reproductive queueing

In the three cooperative breeding scenarios, helpers queue locally in a breeding territory and may inherit the breeder position after the breeder’s death. Helpers are sorted according to their age (from old to young), and a weight for each helper determines how likely a given helper is to inherit the breeder position relative to the other helpers. The weight of the first (and thus oldest) helper is always set to *w*_1_ = 1. The weight of the subsequent helpers is then calculated as

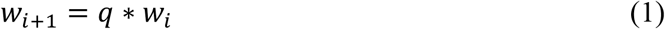

where *q* is a parameter that determines how strongly the probability of a helper to become the new breeder depends on the helper’s age rank among the helpers within the breeding territory. If *q* = 0, then the oldest helper always inherits the breeder position (“strictly age-dependent reproductive queueing”). If 0 < *q* < 1, then relatively older helpers are more likely to inherit the breeder position (“weakly age-dependent reproductive queueing”). If *q* = 1, then the probability to inherit the breeder position is independent of age (“age-independent reproductive queueing”).

### Reproduction

We assume that the number of offspring produced by a breeder increases with the number of helpers in the breeding territory. This is modelled with the diminishing return function

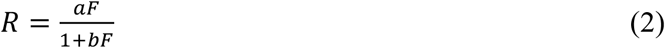

where *F* is the number of females in the breeding territory, *a* is a “maximum productivity”-parameter that determines the maximum number of offspring at large group sizes and *b* is a further parameter that determines the rate at which the maximum is approached. Offspring are produced with even sex ratios. Females mate at independence with one random male. We limited the number of females per breeding territory to *F*_max_ for computational reasons.

### Model analysis

We implemented the model in C++ and compiled it with g++ 4.8.5.. We analysed and visualised model results in R 4.1.0^65^ using the packages *ini*^66^, *gridExtra*^67^, *scales*^68^, *tidyverse*^69^, *cowplot*^70^, *ggpubr*^71^ and *MetBrewer*^72^. We ran simulations until time point *t*end. At this point the simulations had reached an evolutionary quasi-equilibrium at which mean lifespans no longer changed systematically over time (Fig. S6). We calculated evolved lifespans as the sum of the cumulative products of age-specific survival probabilities (“life expectancy”). In order to obtain estimates for relatedness between breeders and helpers, we gave individuals a selectively neutral gene locus that was randomly mutated without mutation bias. We then used the mean of the two homologous loci from all breeders and one random helper per breeding territory to calculate relatedness within breeding territories as the covariance between the breeder and helper gene values divided by the product of the standard deviations of the breeder and helper gene values.

## Acknowledgements

We thank Thijs Janzen for helping us to automatize simulations. We are grateful to Chris Faulkes, Dario Josi, Charli Davies and David Baracchi for giving us permission to use their photographs. We would like to thank the Center for Information Technology of the University of Groningen for providing access to the Peregrine high performance computing cluster. JJK was supported by an Adaptive Life grant by the University of Groningen.

## Author contributions

JJK and IP conceptualised and implemented the model. JJK and BHK analysed the model. All authors contributed to the writing of the manuscript.

## Literature

1. Kreider, J. J., Pen, I. & Kramer, B. H. Antagonistic pleiotropy and the evolution of extraordinary lifespans in eusocial organisms. Evolution Letters 5, 178–186 (2021).

2. Kramer, B. H., van Doorn, S. G., Arani, B. M. S. & Pen, I. Eusociality and the evolution of aging in superorganisms. bioRxiv (2021) doi:10.1101/2021.05.06.442925.

3. Pen, I. & Flatt, T. Asymmetry, division of labour and the evolution of ageing in multicellular organisms. Phil. Trans. R. Soc. B 376, 20190729 (2021).

4. Arnold, K. E. & Owens, I. P. F. Cooperative breeding in birds: a comparative test of the life history hypothesis. Proc. R. Soc. B 265, 739–745 (1998).

5. Downing, P. A., Cornwallis, C. K. & Griffin, A. S. Sex, long life and the evolutionary transition to cooperative breeding in birds. Proc. R. Soc. B 282, 20151663 (2015).

6. Williams, S. A. & Shattuck, M. R. Ecology, longevity and naked mole-rats: confounding effects of sociality? Proc. R. Soc. B. 282, 20141664 (2015).

7. Healy, K. Eusociality but not fossoriality drives longevity in small mammals. Proc. R. Soc. B. 282, 20142917 (2015).

8. Lukas, D. & Clutton-Brock, T. Life histories and the evolution of cooperative breeding in mammals. Proc. R. Soc. B. 279, 4065–4070 (2012).

9. Thorley, J. The case for extended lifespan in cooperatively breeding mammals: a re-appraisal. PeerJ 8, e9214 (2020).

10. Keller, L. & Genoud, M. Extraordinary lifespans in ants: a test of evolutionary theories of ageing. Nature 389, 958–960 (1997).

11. Hatchwell, B. J. & Komdeur, J. Ecological constraints, life history traits and the evolution of cooperative breeding. Animal Behaviour 59, 1079–1086 (2000).

12. Pen, I. & Weissing, F. J. Towards a unified theory of cooperative breeding: the role of ecology and life history re-examined. Proc. R. Soc. Lond. B 267, 2411–2418 (2000).

13. Kokko, H. & Lundberg, P. Dispersal, migration, and offspring retention in saturated habitats. The American Naturalist 157, 188–202 (2001).

14. Kokko, H. & Ekman, J. Delayed dispersal as a route to breeding: territorial inheritance, safe havens, and ecological constraints. The American Naturalist 160, 468–484 (2002).

15. Hamilton, W. D. The genetical evolution of social behaviour. I. Journal of Theoretical Biology 7, 1–16 (1964).

16. Hamilton, W. D. The genetical evolution of social behaviour. II. Journal of Theoretical Biology 7, 17–52 (1964).

17. Hatchwell, B. J. The evolution of cooperative breeding in birds: kinship, dispersal and life history. Phil. Trans. R. Soc. B 364, 3217–3227 (2009).

18. Cornwallis, C. K., West, S. A., Davis, K. E. & Griffin, A. S. Promiscuity and the evolutionary transition to complex societies. Nature 466, 969–972 (2010).

19. Lukas, D. & Clutton-Brock, T. Cooperative breeding and monogamy in mammalian societies. Proc. R. Soc. B. 279, 2151–2156 (2012).

20. Leadbeater, E., Carruthers, J. M., Green, J. P., Rosser, N. S. & Field, J. Nest inheritance is the missing source of direct fitness in a primitively eusocial insect. Science 333, 874–876 (2011).

21. Zöttl, M., Heg, D., Chervet, N. & Taborsky, M. Kinship reduces alloparental care in cooperative cichlids where helpers pay-to-stay. Nat Commun 4, 1341 (2013).

22. Quiñones, A. E., van Doorn, G. S., Pen, I., Weissing, F. J. & Taborsky, M. Negotiation and appeasement can be more effective drivers of sociality than kin selection. Phil. Trans. R. Soc. B 371, 20150089 (2016).

23. Kingma, S. A. Direct benefits explain interspecific variation in helping behaviour among cooperatively breeding birds. Nat Commun 8, 1094 (2017).

24. Hamilton, W. D. The moulding of senescence by natural selection. Journal of Theoretical Biology 12, 12–45 (1966).

25. Kramer, B. H., van Doorn, G. S., Weissing, F. J. & Pen, I. Lifespan divergence between social insect castes: challenges and opportunities for evolutionary theories of aging. Current Opinion in Insect Science 16, 76–80 (2016).

26. Medawar, P. B. An unsolved problem of biology: an inaugural lecture delivered at university college, London, 6 December, 1951. (H.K. Lewis and Company, 1952).

27. Sherman, P. W. & Jarvis, J. U. M. Extraordinary life spans of naked mole-rats (*Heterocephalus glaber*). Journal of Zoology 258, 307–311 (2002).

28. O’Riain, M. J. & Faulkes, C. G. African mole-rats: eusociality, relatedness and ecological constraints. in Ecology of social evolution 207–223 (Springer, 2008).

29. Clarke, F. M. & Faulkes, C. G. Dominance and queen succession in captive colonies of the eusocial naked mole–rat, *Heterocephalus glaber*. Proc. R. Soc. Lond. B 264, 993–1000 (1997).

30. Van der Westhuizen, L., Jarvis, J. U. & Bennett, N. C. A case of natural queen succession in a captive colony of naked mole-rats, Heterocephalus glaber. African Zoology 48, 56–63 (2013).

31. Wong, M. & Balshine, S. The evolution of cooperative breeding in the African cichlid fish, Neolamprologus pulcher. Biological Reviews 86, 511–530 (2011).

32. Stiver, K. A., Dierkes, P., Taborsky, M., Lisle Gibbs, H. & Balshine, S. Relatedness and helping in fish: examining the theoretical predictions. Proc. R. Soc. B. 272, 1593–1599 (2005).

33. Dierkes, P., Heg, D., Taborsky, M., Skubic, E. & Achmann, R. Genetic relatedness in groups is sex‐specific and declines with age of helpers in a cooperatively breeding cichlid. Ecology Letters 8, 968–975 (2005).

34. Richardson, D. S., Burke, T. & Komdeur, J. Direct benefits and the evolution of female-biased cooperative breeding in Seychelles warblers. Evolution 56, 2313–2321 (2002).

35. Eikenaar, C., Richardson, D., Brouwer, L. & Komdeur, J. Parent presence, delayed dispersal, and territory acquisition in the Seychelles warbler. Behavioral Ecology 18, 874–879 (2007).

36. Groenewoud, F. et al. Subordinate females in the cooperatively breeding Seychelles warbler obtain direct benefits by joining unrelated groups. J Anim Ecol 87, 1251–1263 (2018).

37. Sumner, S., Casiraghi, M., Foster, W. & Field, J. High reproductive skew in tropical hover wasps. Proc. R. Soc. Lond. B 269, 179–186 (2002).

38. Bridge, C. & Field, J. Queuing for dominance: gerontocracy and queue-jumping in the hover wasp Liostenogaster flavolineata. Behav Ecol Sociobiol 61, 1253–1259 (2007).

39. Shreeves, G. & Field, J. Group size and direct fitness in social queues. The American Naturalist 159, 81–95 (2002).

40. Cronin, A. & Field, J. Social aggression in an age-dependent dominance hierarchy. Behav 144, 753–765 (2007).

41. Brouwer, L., Richardson, D. S. & Komdeur, J. Helpers at the nest improve late-life offspring performance: evidence from a long-term study and a cross-foster experiment. PLoS ONE 7, e33167 (2012).

42. Canestrari, D., Marcos, J. M. & Baglione, V. Helpers at the nest compensate for reduced maternal investment in egg size in carrion crows. Journal of Evolutionary Biology 24, 1870–1878 (2011).

43. Doerr, E. D. & Doerr, V. A. J. Positive effects of helpers on reproductive success in the brown treecreeper and the general importance of future benefits. J Anim Ecology 76, 966–976 (2007).

44. Hatchwell, B. J. Helpers increase long-term but not short-term productivity in cooperatively breeding long-tailed tits. Behavioral Ecology 15, 1–10 (2004).

45. Koenig, W. D., Walters, E. L. & Barve, S. Does helping-at-the-nest help? The case of the acorn woodpecker. Front. Ecol. Evol. 7, 272 (2019).

46. Preston, S. A. J., Briskie, J. V. & Hatchwell, B. J. Adult helpers increase the recruitment of closely related offspring in the cooperatively breeding rifleman. Behavioral Ecology 27, 1617–1626 (2016).

47. Schwarz, M. P. Local resource enhancement and sex ratios in a primitively social bee. Nature 331, 346–348 (1988).

48. Kramer, B. H., Scharf, I. & Foitzik, S. The role of per-capita productivity in the evolution of small colony sizes in ants. Behavioral Ecology and Sociobiology 68, 41–53 (2014).

49. Carey, J. R. Demographic mechanisms for the evolution of long life in social insects. Experimental Gerontology 36, 713–722 (2001).

50. Carey, J. R. & Judge, D. S. Life span extension in humans is self-reinforcing: a general theory of longevity. Population and Development Review 27, 411–436 (2001).

51. Korb, J. & Heinze, J. Ageing and sociality: why, when and how does sociality change ageing patterns? Phil. Trans. R. Soc. B 376, 20190727 (2021).

52. Hammers, M. et al. Breeders that receive help age more slowly in a cooperatively breeding bird. Nat Commun 10, 1301 (2019).

53. Downing, P. A., Griffin, A. S. & Cornwallis, C. K. Hard-working helpers contribute to long breeder lifespans in cooperative birds. Phil. Trans. R. Soc. B 376, 20190742 (2021).

54. Hammers, M. et al. Helpers compensate for age‐related declines in parental care and offspring survival in a cooperatively breeding bird. Evolution Letters 5, 143–153 (2021).

55. Kramer, B. H. & Schaible, R. Colony size explains the lifespan differences between queens and workers in eusocial Hymenoptera. Biological Journal of the Linnean Society 109, 710–724 (2013).

56. West, S. A., Pen, I. & Griffin, A. S. Cooperation and competition between relatives. Science 296, 72–75 (2002).

57. Lee, R. D. Rethinking the evolutionary theory of aging: transfers, not births, shape senescence in social species. Proceedings of the National Academy of Sciences 100, 9637–9642 (2003).

58. Bourke, A. F. G. Kin selection and the evolutionary theory of aging. Annual Review of Ecology, Evolution, and Systematics 3, 103–128 (2007).

59. Lee, R. Sociality, selection, and survival: simulated evolution of mortality with intergenerational transfers and food sharing. Proceedings of the National Academy of Sciences 105, 7124–7128 (2008).

60. Croft, D. P., Brent, L. J. N., Franks, D. W. & Cant, M. A. The evolution of prolonged life after reproduction. Trends in Ecology & Evolution 30, 407–416 (2015).

61. Schmidt, L., Sobotka, T., Bentzen, J. G., Nyboe Andersen, A., & on behalf of the ESHRE Reproduction and Society Task Force. Demographic and medical consequences of the postponement of parenthood. Human Reproduction Update 18, 29–43 (2012).

62. Wyshak, G. & Frisch, R. E. Evidence for a secular trend in age of menarche. New England journal of medicine 306, 1033–1035 (1982).

63. Vaupel, J. W. Biodemography of human ageing. Nature 464, 536–542 (2010).

64. Burger, O., Baudisch, A. & Vaupel, J. W. Human mortality improvement in evolutionary context. Proceedings of the National Academy of Sciences 109, 18210–18214 (2012).

65. R Core Team. R: A Language and Environment for Statistical Computing. (R Foundation for Statistical Computing, 2021).

66. Dias, D. V. ini: read and write ‘.ini’ files. (https://CRAN.R-project.org/package=ini, 2018).

67. Auguie, B. gridExtra: miscellaneous functions for ‘Grid’ graphics. (https://CRAN.R-project.org/package=gridExtra, 2017).

68. Wickham, H. & Seidel, D. scales: scale functions for visualization. (https://CRAN.R-project.org/package=scales, 2020).

69. Wickham, H. et al. Welcome to the tidyverse. Journal of Open Source Software 4, 1686 (2019).

70. Wilke, C. O. cowplot: streamlined plot theme and plot annotations for ‘ggplot2’. (https://CRAN.R-project.org/package=cowplot, 2019).

71. Kassambara, A. ggpubr: ‘ggplot2’ based publication ready plots. (https://CRAN.R-project.org/package=ggpubr, 2020).

72. Mills, B. R. MetBrewer: color palettes inspired by works at the Metropolitan Museum of Art. (2021).

